# HSC70 regulates cold-induced caspase-1 hyperactivation by an autoinflammation-causing mutant of cytoplasmic immune receptor NLRC4

**DOI:** 10.1101/578138

**Authors:** Akhouri Kishore Raghawan, Rajashree Ramaswamy, Vegesna Radha, Ghanshyam Swarup

**Affiliations:** CSIR-Centre for Cellular and Molecular Biology, Hyderabad-500007, India

**Keywords:** NLRC4, HSC70, caspase-1, H443P mutant, inflammasome, cold hypersensitivity

## Abstract

NLRC4 is an innate immune receptor, which upon detection of certain pathogens or internal distress signal, initiates caspase-1 mediated inflammatory response. A gain-of-function mutation, H443P in NLRC4, causes familial cold autoinflammatory syndrome (FCAS) characterized by cold-induced hyperactivation of caspase-1 and inflammation. Here, we show that heat shock cognate protein 70 (HSC70) complexes with NLRC4 and negatively regulates caspase-1 activation by NLRC4-H443P. Compared to NLRC4, the structurally altered NLRC4-H443P shows enhanced interaction with HSC70. Knockdown of HSC70 or inhibition of its ATPase activity enhances caspase-1 activation by NLRC4-H443P. Exposure to subnormal temperature resulted in reduced interaction of NLRC4-H443P with HSC70, and an increase in its ability to form ASC-specks and activate caspase-1. By demonstrating that HSC70 differentially interacts with NLRC4-H443P mutant in a temperature-dependent manner to regulate caspase-1 activation, we provide a mechanism for cold-induced inflammation seen in FCAS patients with NLRC4-H443P mutation.

## Introduction

Cytoplasmic and membrane-bound receptors mediate innate immune response upon sensing invading pathogens or intracellular danger signals^1–3^. NLRC4 is a member of Nod-like receptor-family of cytoplasmic immune receptors. It is expressed in immune cells such as monocytes, macrophages and neutrophils. In addition, NLRC4 functions are also known in non-hematopoietic cells such as lung and intestinal epithelial cells and brain cells^4–7^. NLRC4 comprises an N-terminal caspase recruitment domain (CARD), a central nucleotide binding and oligomerisation domain (NBD), a winged helix domain (WHD), two helical domains HD1 and HD2, and a C-terminal leucine-rich repeat (LRR) domain^8–10^. NLRC4 is activated upon detection of bacterial flagellin and rod proteins to form a multimolecular complex known as inflammasome, and activates caspase-1. NLRC4 can engage and activate caspase-1 either directly through its CARD domain or through an adapter protein, apoptosis-associated speck-like protein containing a CARD (ASC)^11^. Activated caspase-1 proteolytically cleaves pro-interleukin (IL)-1β and pro-IL-18 into mature cytokines that mediate an inflammatory response downstream of NLRC4^11–17^. NLRC4 is maintained in an ADP-bound auto-inhibited state through intermolecular interactions between NBD and WHD of NLRC4, and ADP, and the LRR domain folds over to inhibit its oligomerisation and activation^8^. Molecules that interact with NLRC4 are likely to alter its configuration, and its ability to modulate downstream signalling events, but these mechanisms are not fully understood.

Mutations in NLRC4^18–24^ and some other cytoplasmic immune receptors like NLRP1^11, 25– 27^, NLRP3^28–31^ and NLRP12^32–34^ cause autoinflammatory syndromes, which occur in the absence of any infection or autoimmunity. One of the mutants of NLRC4, H443P causes familial cold autoinflammatory syndrome (FCAS) in heterozygous individuals^18^. FCAS is a mild form of autoinflammatory syndrome characterized by arthralgia, intermittent fever and skin rashes upon exposure of the individual to subnormal temperatures. NLRC4-H443P undergoes auto-oligomerisation leading to constitutive activation of caspase-1 and maturation of cytokine IL-1β. Hyper-inflammatory response is seen in transgenic mice expressing NLRC4-H443P upon exposure to cold water at 4ºC^18^. This inflammation in NLRC4-H443P transgenic mice is due to caspase-1 mediated IL-1β maturation. Overexpression of NLRC4-H443P in HEK293T cells causes increased activation of caspase-1 when exposed to temperature of 32ºC^18^. NLRC4-H443P differs from the wild type (WT) NLRC4 in properties like altered interaction with SUG1, a proteasomal component, higher ubiquitination and increased interaction with ubiquitinated cellular proteins. In addition to constitutive caspase-1 activation and inflammation, NLRC4-H443P induces SUG1 and FADD-dependent caspase-8 activation and apoptosis^35^.

Heat shock proteins (HSPs) are a family of molecular chaperones involved in protein-homeostasis in a cell and are broadly conserved across all vertebrates^36^. HSPs are involved in physiological processes like folding of newly synthesized polypeptides, re-folding or degradation of misfolded proteins, protein sorting, transport and secretion, autophagy, apoptosis, stress response during hypothermia/hyperthermia, and inflammatory immune response^37–44^. HSPs have been divided into two broad categories based on their size and mode of function. Larger HSPs i.e. HSP70, HSC70, GRP78 and HSP90 are 70-90KDa proteins and require ATPase activity for most of their functions while smaller HSPs i.e. HSP27 and HSP40 mostly act as co-chaperones to larger HSPs and do not possess ATPase activity. Many HSPs are induced during stress response. Heat shock cognate protein 70 (HSC70), encoded by HSPA8 gene in humans, is a constitutively and ubiquitously expressed member of HSP-family and constitutes 1-2% of total cellular proteins. Unlike many HSPs, HSC70 is not induced in response to higher temperature. HSC70 possesses an N-terminal ATPase domain, a substrate binding domain (SBD) and a C-terminal lid domain^45^. It is involved in promoting protein folding and in processing of mis-folded proteins^37, 38^. HSC70, in collaboration with other co-chaperones and HSP90 is involved in maintaining protein homeostasis by mediating proteasomal degradation of misfolded proteins^39,40, 46, 47^.

The chaperone function of HSC70 helps in proper folding of newly synthesized proteins and this function involves transient interactions with short hydrophobic (or hydrophobic-basic) sequences in the partially folded / unfolded proteins. These interactions of HSC70 increase with increasing temperature in the physiological range of 30-37°C. HSC70 undergoes a temperature-dependent reversible conformational change beginning at about 30°C, which increases its interaction with peptides and unfolded proteins, and its chaperoning activity^36, 48^. Thus, HSC70 exists in active and inactive states, and the proportion of active pool is determined by temperature-dependent conformational change. These properties of HSC70 indicate that it is a potential candidate to modulate temperature-dependent functional properties of client proteins in the physiological temperature range of 30-37°C.

The molecular mechanism of inflammation triggered by exposure to subnormal temperature is by and large not known. In this study, we attempted to understand how caspase-1 hyper-activation and inflammation occurs downstream of FCAS-causing NLRC4-H443P mutant. We have identified HSC70 and HSP70 as novel interacting partners of NLRC4 and show that HSC70 negatively regulates the inflammasome function of NLRC4-H443P in an ATPase-dependent manner. Upon exposure to subnormal temperatures, NLRC4-H443P interaction with cellular HSC70 decreases, resulting in increased inflammasome formation and caspase-1 activation. Our results suggest that HSC70 plays an important role in modulating temperature sensitive properties of NLRC4-H443P.

## Results

### HSC70 forms a complex with NLRC4 and shows enhanced interaction with H443P mutant

Gain-of-function mutations in NLRC4 cause auto-inflammatory disorders in humans^24^. Fig.1A shows a schematic indicating positions of various disease associated mutations in NLRC4 protein. A missense mutation, H443P in NLRC4 causes cold hypersensitivity in heterozygous individuals. While studying differential interaction of SUG1 with WT-NLRC4 and NLRC4-H443P by immunoprecipitation,^35^ we observed a prominent 70KDa cellular polypeptide in NLRC4 immunoprecipitates from cells transiently expressing GFP-fusion proteins of WT-NLRC4 or NLRC4-H443P (Supplementary Figure 1A). The prominence of this protein suggested that it could be one of the HSPs of 70KDa, which play an important role in maintaining cellular homeostasis in response to abnormal temperatures. Western blot analysis of immunoprecipitates showed that cellular HSC70 forms a complex with NLRC4 (Fig. 1B). Higher levels of HSC70 were seen repeatedly in immunoprecipitates of H443P mutant compared to that of WT-NLRC4, suggesting that HSC70 complexes with H443P mutant with higher affinity than with WT-NLRC4 (Fig. 1B, C). We examined interaction of HSC70 with two other disease-associated mutants of NLRC4, T337S and V341A, which also cause constitutive caspase-1 activation. T337S and V341A mutants of NLRC4 do not cause FCAS but are involved in causing other autoinflammatory syndromes^19, 20^. Interestingly, compared to WT-NLRC4, HSC70 showed stronger interaction only with the temperature sensitive mutant NLRC4-H443P, but not with NLRC4-T337S or NLRC4-V341A (Fig. 1B). Interaction of endogenous NLRC4 with HSC70 was seen in differentiated THP1 cells by immunoprecipitation indicating that complex formation was not due to forced expression of the proteins (Fig. 1D).

**Fig. 1:**
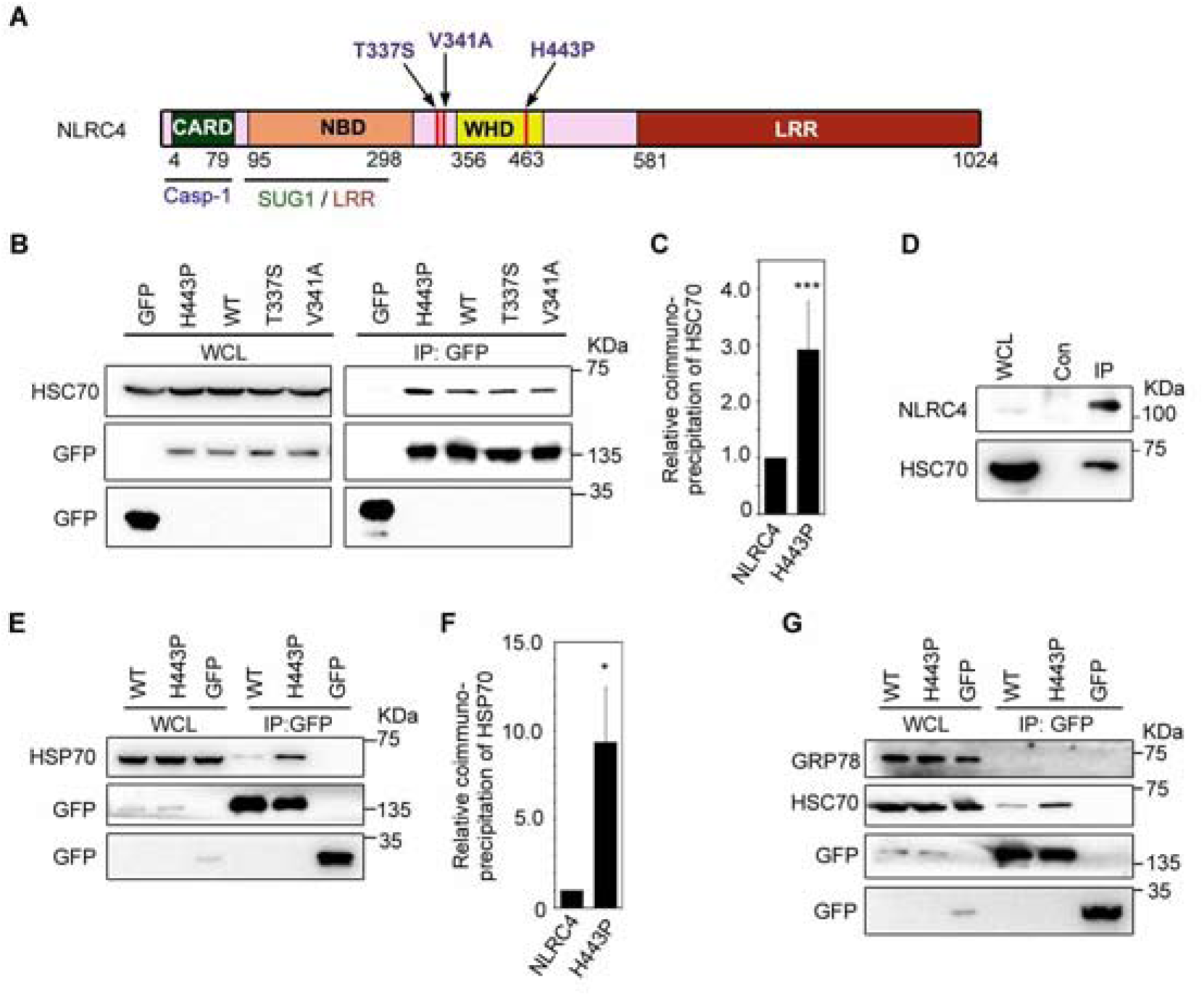
HSC70 and HSP70 are present in cellular complexes formed by NLRC4. **(A)** Schematic showing domain organization of NLRC4. Sites of disease associated mutations in NLRC4 are indicated. **(B)** HSC70 is present in the complex formed by NLRC4 and shows enhanced interaction with NLRC4-H443P mutant. GFP-tagged NLRC4 and its mutant were transiently expressed in HEK293T cells and lysates were subjected to immunoprecipitation using GFP antibody. Western blot analysis showed presence of HSC70 in the immunoprecipitates (IP) of WT-NLRC4 as well as H443P, T337S and V341A mutants. WCL, whole cell lysates **(C)** Bar diagram shows quantitation of relative abundance of HSC70 in immunoprecipitates of H443P mutant compared to WT-NLRC4 from 8 independent experiments; n=8. *** p<0.0005. **(D)** Western blot shows interaction between endogenous NLRC4 and HSC70 in differentiated THP1 cells. Cells were lysed and subjected to immunoprecipitation using NLRC4 antibody (IP) or normal IgG (Con) and analysed by western blotting. **(E)** HSP70 forms a complex with NLRC4 and shows enhanced interaction with NLRC4-H443P. Lysates of cells transfected with indicated constructs were subjected to immunoprecipitation and western blotting. **(F)** Bar diagram shows relative abundance of HSP70 in immunoprecipitates of WT-NLRC4 and NLRC4-H443P; n=3. * p<0.05. **(G)** GRP78is not present in the cellular complexes formed by NLRC4.

In addition to HSC70, HSP70 was detected in the immunoprecipitates obtained from lysates of HEK293T cells transiently expressing WT-NLRC4 or NLRC4-H443P. Higher levels of HSP70 were seen in complexes with NLRC4-H443P compared to that of WT-NLRC4 (Fig. 1E, F). GRP78, another HSP70 member, which is localized to the endoplasmic reticulum and is required for folding of newly synthesized proteins, was not detected in NLRC4 or NLRC4-H443P immunoprecipitates under similar experimental conditions (Fig. 1G). These results identified HSC70 and HSP70 as two novel interacting partners of NLRC4 and suggested a possible role for HSC70 / HSP70 in signal transduction downstream of the temperature sensitive mutant NLRC4-H443P.

### Domain requirements for interaction between NLRC4 and HSC70

In order to identify the domains in NLRC4 required for its interaction with HSC70, we transfected deletion constructs expressing GFP-fusion proteins of various domains of NLRC4 (Fig. 2A) in HEK293T cells and subjected the lysates to immunoprecipitation using GFP-antibody. HSC70 was present in the immunoprecipitates of deletion constructs expressing ΔLRR-NLRC4 (which lacks LRR domain) as well as in ΔCARD-ΔLRR-NLRC4, which lacks both LRR domain and CARD (Fig. 2B). GFP-LRR and GFP-aa91-253 interacted with endogenous HSC70, but GFP-CARD failed to coprecipitate HSC70 in our assays (Fig. 2C). These results indicated that NBD and LRR domain of NLRC4 can independently interact with HSC70, and CARD domain is neither sufficient nor essential for interaction. In-vitro GST pulldown assays using GST-HSC70 showed that GFP-LRR and GFP-aa91-253, but not GFP can interact with HSC70 (Fig. 2D). For GST pulldown assays, binding reaction was carried out at 37°C because no interaction of GST-HSC70 with NLRC4 or NLRC4-H443P was seen at 4°C, a condition generally used for these assays (data not shown). HSC70 is known to bind to its substrates primarily through its carboxyl-terminal substrate binding domain (aa394-546) (SBD) (shown schematically, Fig. 2A), which has affinity for extended hydrophobic motifs in the target protein^49, 50^. GST-SBD interacted with NLRC4 and NLRC4-H443P in in-vitro binding assays (Fig. 2E).

**Fig. 2:**
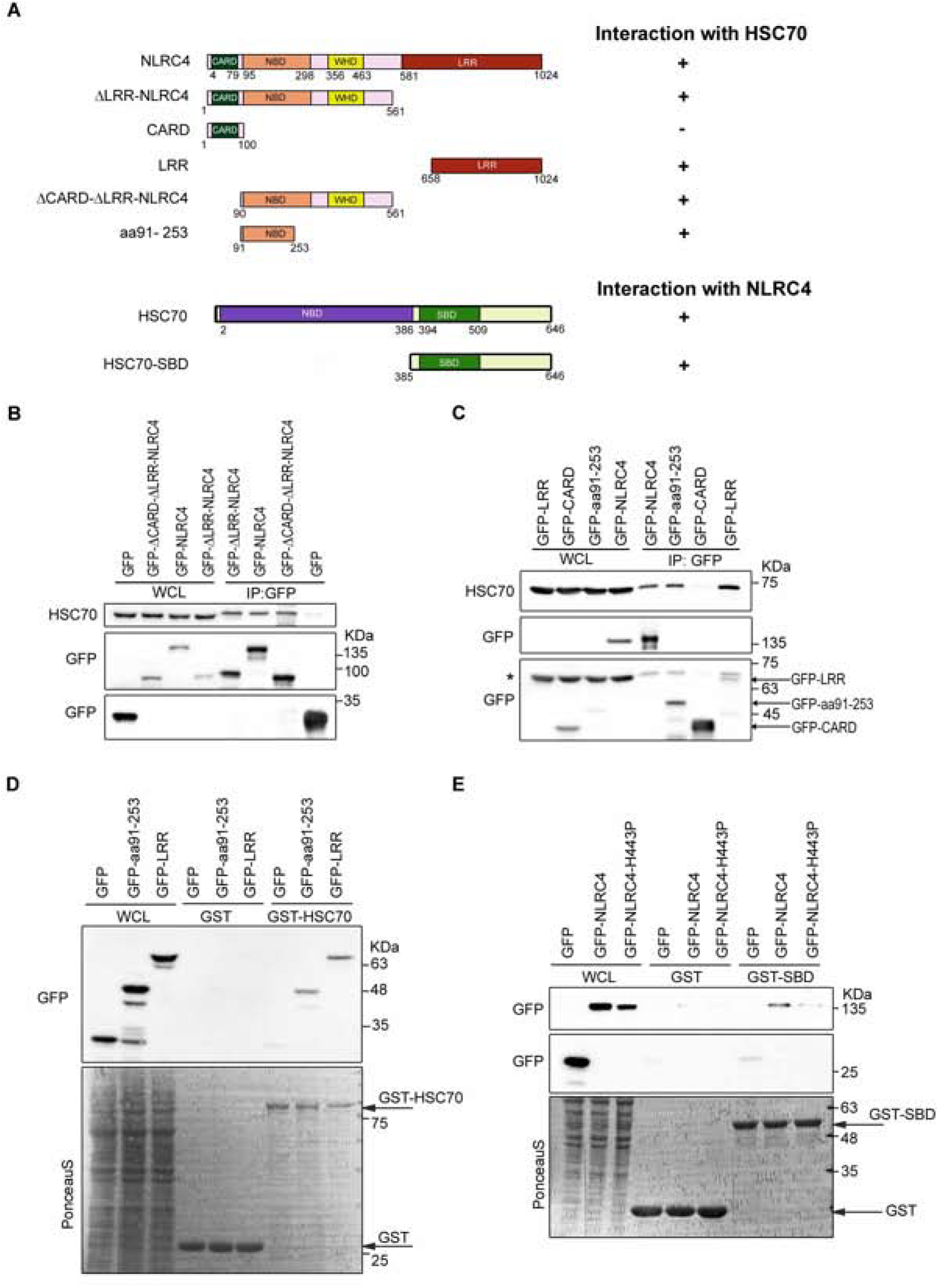
Identification of domains in NLRC4 and HSC70 required for complex formation. **(A)** Schematic showing various deletion constructs of NLRC4 and HSC70. + in the right hand side indicates positivity for indicated interaction. **(B and C)** NBD (aa91-253) and LRR (aa658-1024) can independently bind to HSC70. GFP-tagged NLRC4 or its deletion constructs were transiently transfected in HEK293T cells for 16h. Whole cell lysates were subjected to immunoprecipitation using GFP antibody and analysed by western blotting to test presence of endogenous HSC70. The blot in (C) was probed with HSC70 antibody followed by reprobing with GFP antibody; * indicates residual signal from bands of endogenous HSC70; arrows indicate bands corresponding to the indicated NLRC4 proteins. **(D)** GST pulldown assay showing binding of GST-HSC70 with GFP-aa91-253 and GFP-LRR. Purified GST or GST-HSC70 were incubated with lysates of HEK293T cells expressing GFP, GFP-aa91-253 or GFP-LRR and bound proteins analysed by western blotting. Arrows point to positions of GST or GST-HSC70 in the PonceauS stained blot. **(E)** Substrate binding domain of HSC70 (SBD) is sufficient to bind with NLRC4 or NLRC4-H443P. GST fusion protein of SBD was incubated with lysates of HEK293T cells expressing WT-NLRC4 or NLRC4-H443Pand bound proteins analysed by western blotting. Arrows indicate expression of the recombinant proteins.

### Ubiquitination of NLRC4-H443P enhances interaction with HSC70

While higher amount of HSC70 was present in cellular complexes formed by NLRC4-H443P compared to WT-NLRC4 (Fig. 1B), no difference was seen in interaction when in-vitro binding assays were carried out (Fig. 3A). We hypothesized that NLRC4-H443P mutant may be undergoing a post-translational modification in cells which is lost or reduced in cell lysates. Previously, we have reported that NLRC4 undergoes ubiquitination and that higher level of ubiquitination is seen on NLRC4-H443P^35^ (Supplementary Figure 1B). Therefore, we carried out cell lysis and GST-pulldown assays in buffers containing N-ethylmaleimide (NEM), an inhibitor of cellular deubiquitinases. Higher amount of NLRC4 as well as NLRC4-H443P complexed with GST-HSC70 when NEM was included in the cell lysis and binding buffers, compared to experiments without NEM (Fig. 3B). Compared to WT-NLRC4, NLRC4-H443P showed enhanced interaction with GST-HSC70 in the presence of NEM (Fig. 3C). De-probing the blot followed by re-probing with ubiquitin antibody, confirmed enhanced ubiquitination on polypeptide corresponding to NLRC4-H443P, compared to WT-NLRC4 in samples processed with NEM (Fig. 3B). These results suggest that HSC70 preferentially interacts with ubiquitinated NLRC4-H443P mutant and enhanced binding of this mutant is likely due to higher ubiquitination of NLRC4-H443P mutant compared to that of WT-NLRC4.

**Fig. 3:**
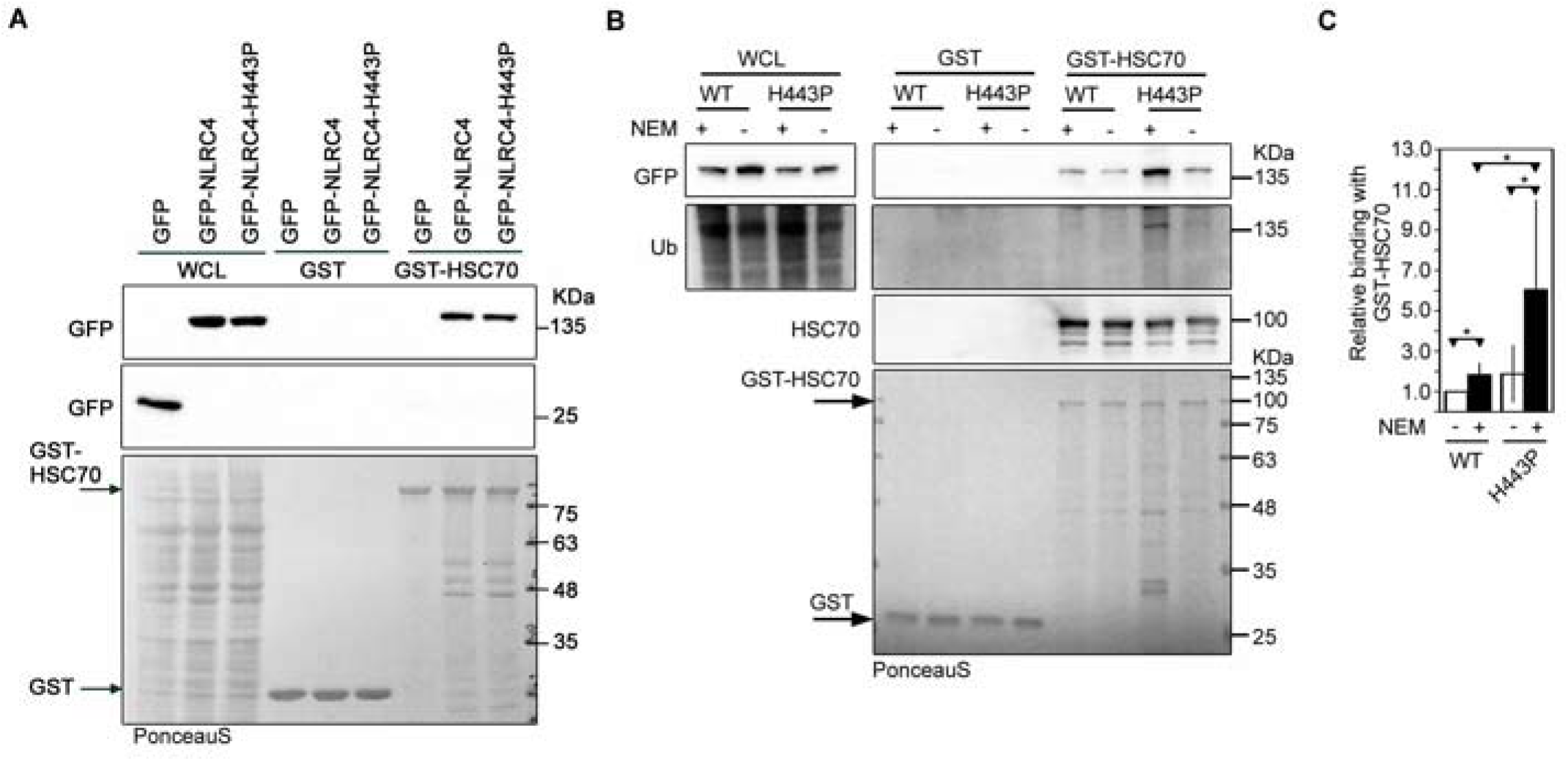
Interaction of HSC70 with NLRC4-H443P is dependent on ubiquitination of NLRC4-H443P. **(A)** GST-HSC70 was incubated with lysates of HEK293T expressing GFP, GFP-NLRC4 or GFP-NLRC4-H443P and bound proteins analysed by western blotting. GST-HSC70 did not show increased binding with NLRC4-H443P compared to WT-NLRC4. **(B)** Lysates of HEK293T expressing GFP-NLRC4 or GFP-NLRC4-H443P were prepared in buffer with or without NEM (10mM) and incubated with GST-HSC70. Bound proteins were analysed by western blotting with indicated antibodies. **(C)** Bar diagram shows relative abundance of NLRC4 or NLRC4-H443P in pulldown samples of GST-HSC70 in the presence or absence of NEM. n=6. **p<0.005 and ***p<0.0005.

### HSC70 negatively regulates caspase-1 activation by NLRC4-H443P

The FCAS-causing NLRC4-H443P constitutively activates caspase-1 leading to pro-inflammatory cytokine IL-1β maturation. To find out if HSC70 plays a role in caspase-1 activation by NLRC4-H443P, we examined the consequence of knockdown of HSC70 using short interfering RNAs (siRNAs). We observed increased caspase-1 activation by NLRC4-H443P compared to WT-NLRC4 in control siRNA transfected samples (Fig. 4A, B). Caspase-1 activation by NLRC4-H443P further increased upon HSC70 knockdown suggesting that HSC70 negatively regulates caspase-1 activation downstream of NLRC4-H443P. There was only a marginal effect of HSC70-knockdown on caspase-1 activation by WT-NLRC4 (Fig. 4A, B). We confirmed the efficacy of siRNA mediated HSC70-knockdown by western blot (Fig. 4C). HSP70 levels remained unaffected in these samples indicating the specificity of siRNA for HSC70 (Fig. 4D).

**Fig. 4:**
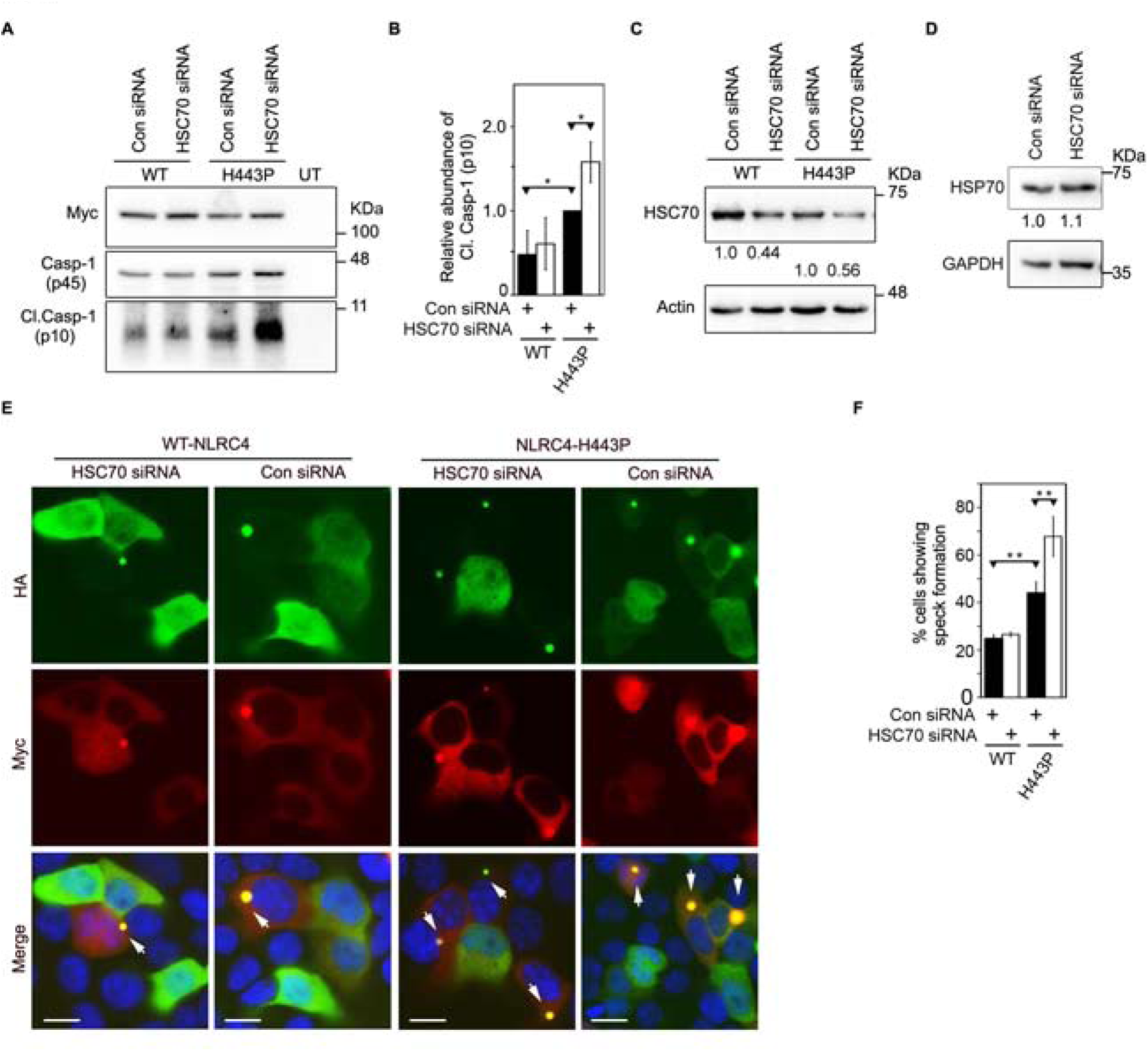
HSC70 negatively regulates caspase-1 activation downstream of NLRC4-H443P. **(A)** HEK293T cells were transfected with control siRNA or siRNA targeted against HSC70 along with caspase-1 and Myc-tagged NLRC4 or NLRC4-H443P. Whole cell lysates were analysed by western blotting for presence of cleaved caspase-1 (Cl.Casp-1, p10). **(B)** Bar diagram shows quantitation of relative abundance of Cl.Casp-1 in cells expressing NLRC4-H443P; n=3. * p<0.05. **(C)** Western blot shows efficacy of HSC70 knockdown by siRNA. **(D)** HSC70 siRNA does not affect cellular HSP70 indicating target specificity. **(E)** Representative images show effect of siRNA mediated knockdown of HSC70 on speck formation by WT-NLRC4 or NLRC4-H443P mutant. Cells co-expressing HA-ASC and Myc-NLRC4 or Myc-NLRC4-H443P were scored for presence or absence of specks. White arrows indicate ASC-specks formed inside cells. Scale bars, 10µm. **(F)** Quantitation of effect of HSC70 knockdown on ASC-mediated speck formation by NLRC4 or NLRC4-H443P. n=4. ** p<0.005.

NLRC4 coexpression enables ASC to form distinct specks in cells, an indicator of inflammasome assembly, which occurs due to oligomerisation of ASC with NLRC4^20, 51^. NLRC4-H443P coexpression resulted in significantly higher number of cells showing ASC-specks compared to those expressing NLRC4 (Fig. 4E, F). ASC mediated speck formation by NLRC4-H443P increased significantly upon HSC70 knockdown (Fig. 4E, F). Importantly, reduction in levels of HSC70 did not affect speck formation by wild type NLRC4.

### HSC70 requires its ATPase activity to regulate inflammasome function of NLRC4-H443P

HSC70 and HSP70 function through ATP-dependent cycles of substrate binding and release^38^. Apoptozole (Az) is a chemical inhibitor of HSC70/HSP70 ATPase activity^52^, and has been used in biochemical studies involving HSC70 ^52–55^. We tested if HSC70 requires its ATPase function in modulating NLRC4-H443P mediated caspase-1 activation by quantitating IL-1β maturation. HEK293T cells transiently expressing caspase-1 and IL-1β along with WT-NLRC4 or NLRC4-H443P were treated with DMSO, 0.2µM or 0.5µM apoptozole. Expression of NLRC4-H443P increased IL-1β maturation, compared to WT-NLRC4 in cell lysates (Fig. 5A, B). Treatment with apoptozole did not have significant effect on WT-NLRC4 induced IL-1β maturation, but increased levels of mature IL-1β in the lysates of NLRC4-H443P transfected cells (Fig. 5A, B). The level of cleaved caspase-1 increased upon treatment with apoptozole in H443P mutant expressing cells but not in WT-NLRC4 expressing cells (Fig. 5C). We also monitored the effect of inhibiting ATPase activity of HSC70 on inflammasome formation by NLRC4-H443P. HEK293T cells expressing HA-tagged ASC and Myc-tagged NLRC4, NLRC4-H443P or NLRC4-V341A were treated with apoptozole or DMSO and examined for presence of ASC-specks. Apoptozole treatment resulted in a significant increase in frequency of specks in NLRC4-H443P expressing cells while there was no effect on speck formation by WT-NLRC4 or NLRC4-V341A (Fig. 5D, E, F, Supplementary Figure 2). These results suggested that HSC70 is dependent on its ATPase activity to regulate inflammasome formation and caspase-1 activation by NLRC4-H443P.

**Fig. 5:**
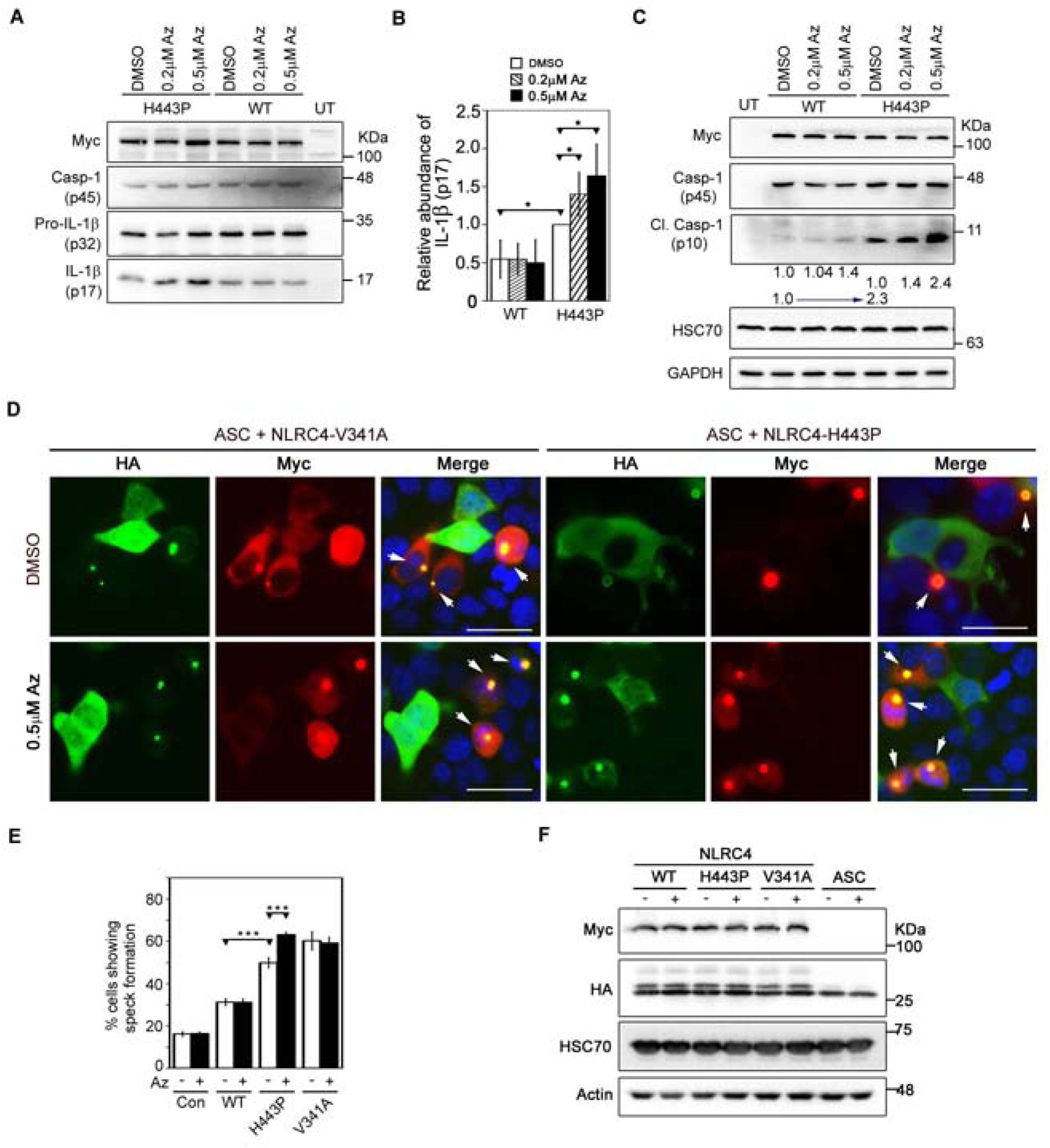
Effect of apoptozole treatment on caspase-1 activation and IL-1β maturation downstream of NLRC4-H443P. **(A)** HEK293T cells were transfected with Myc-tagged NLRC4 or NLRC4-H443P along with caspase-1 and pro-IL-1β. Whole cell lysates were analysed by western blotting for presence of mature IL-1β (p17). Apoptozole was added to the culture medium, in concentrations as indicated. **(B)** Bar diagram shows quantitation of relative levels of mature IL-1β in whole cell lysates upon apoptozole treatment; n=3. * p<0.05. **(C)** Western blot analysis showing increase in caspase-1 activation by NLRC4-H443P upon treatment with apoptozole. Lysates analysed by western blotting for abundance of cleaved caspase-1 (p10). **(D)** Representative immunofluorescence images showing effect of apoptozole treatment (0.5µM; 6h) on speck formation by Myc-NLRC4-V341A and Myc-NLRC4-H443P coexpressed with HA-ASC. White arrows indicate specks. Scale bars, 20µm. **(E)** Quantitation of effect of apoptozole treatment on ASC-speck formation by NLRC4 and its mutants. n=4; *** p<0.0005. **(F)** Expression levels of indicated proteins were checked by western blotting.

### Effect of subnormal temperature on inflammasome formation and caspase-1 activation by NLRC4-H443P

It is known that caspase-1 mediated IL-1β maturation in NLRC4-H443P mutant expressing cells increases upon exposure to subnormal temperature^18^. This was validated by examining the effect of subnormal temperature of 28ºC on ASC-mediated speck formation by H443P mutant, V341A mutant and WT-NLRC4. We observed that NLRC4-H443P and NLRC4-V341A expressing cells showed significantly higher percentage of cells with specks compared to WT-NLRC4 expressing cells. Upon exposure to subnormal temperature of 28ºC, NLRC4-H443P showed further increase in speck formation while there was no significant change in speck formation by WT-NLRC4 or NLRC4-V341A (Fig. 6A, B, Supplementary Figure 3). These results suggest that NLRC4-H443P mutant shows enhanced inflammasome formation upon exposure to lower temperature, whereas V341A mutant does not. As expected, we observed that NLRC4-H443P showed increased caspase-1 activation and IL-1β maturation, and NLRC4-H443P mediated caspase-1 activation and IL-1β maturation further increased upon exposure to subnormal temperature (Fig. 6C, D, E).

**Fig. 6:**
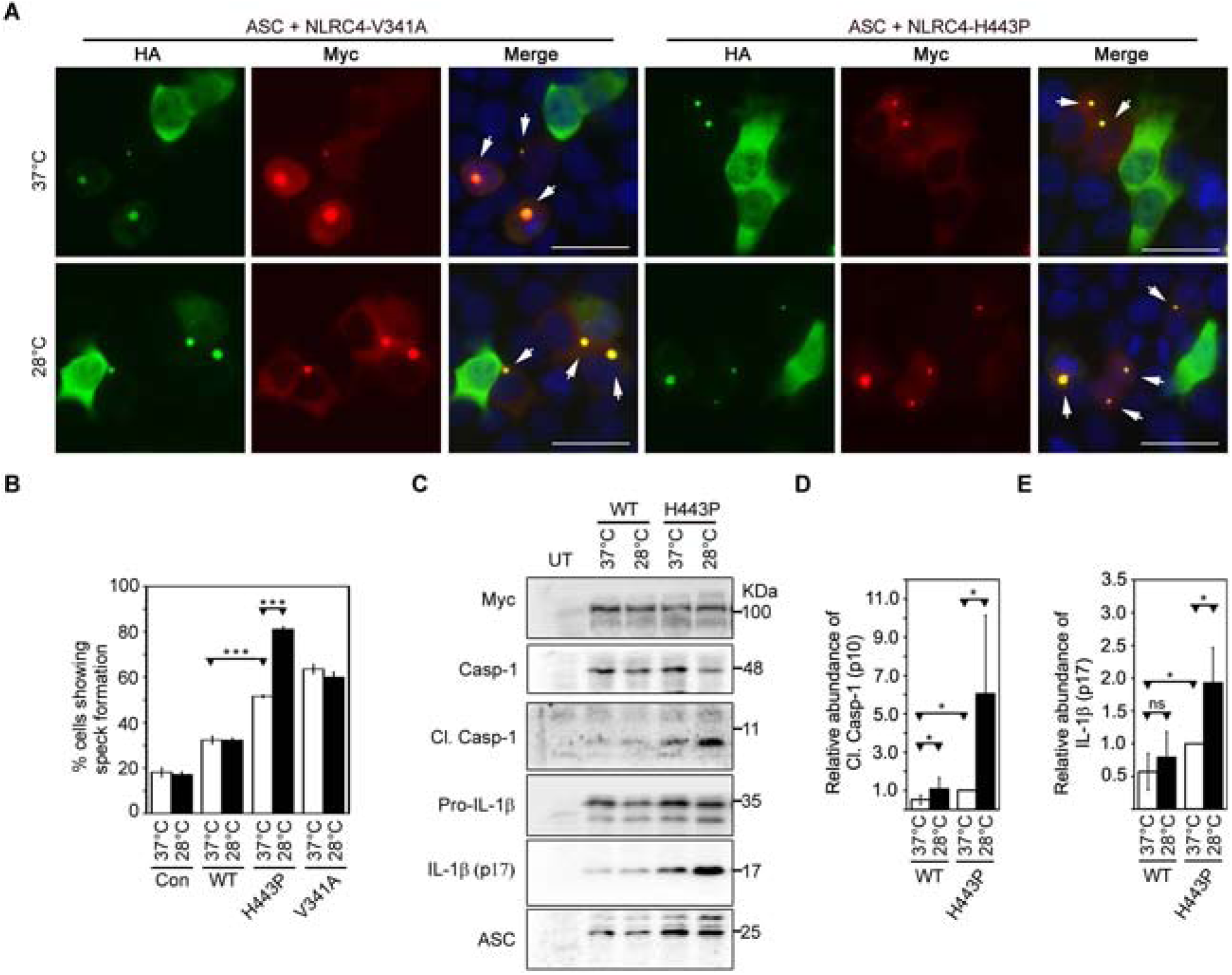
NLRC4-H443P expressing cells exposed to subnormal temperature show increase in inflammasome formation and caspase-1 activation. **(A)** Representative immunofluorescence images show effect of exposure to 28°C for 6h on ASC-speck formation in cells expressing HA-ASC along with NLRC4-V341A or NLRC4-H443P. White arrows indicate specks. DAPI was used to stain nucleus. Scale bars, 20µm. **(B)** Quantitation of ASC-speck formation in response to subnormal temperature. n=6. ** p<0.005 and *** p<0.0005. **(C)** Western blot analysis of lysates of HEK293T cells expressing caspase-1 and HA-ASC along with Myc-NLRC4, Myc-NLRC4-H443P or Myc-NLRC4-V341A. Lysates were analysed for presence of Cl. casp-1 (p10). Bar diagram shows quantitation of effect of exposure to subnormal temperature on caspase-1 activation **(D)** and IL-1β maturation **(E).** n=6. * p<0.05.

### Effect of subnormal temperature on interaction of NLRC4-H443P with HSC70 and ASC

We examined the effect of exposure to subnormal temperature on interaction of HSC70 with NLRC4 or NLRC4-H443P. HEK293T cells expressing Myc-tagged NLRC4 or NLRC4-H443P were maintained at 37ºC or exposed to 28ºC before lysis. Lysates were subjected to immunoprecipitation with Myc antibody and Western blotting. Significantly reduced levels of HSC70 complexed with NLRC4-H443P in cells exposed to subnormal temperature compared to that in cells at 37ºC (Fig. 7A, B, C). The interaction of wild type NLRC4 with HSC70 showed a small reduction upon exposure to lower temperature (Fig. 7A, B). In-vitro binding assays also showed that GST-HSC70 binds weakly with NLRC4-H443P at 28ºC compared to that at 37ºC (Fig. 7D, E). Significant effect of subnormal temperature on binding of GST-HSC70 with wild type NLRC4 was seen (Fig. 7D, E). Western blot analysis of immunoprecipitates from lysates of cells expressing ASC along with NLRC4-H443P showed that NLRC4-H443P was more abundant in cellular complexes formed by ASC in cells exposed to subnormal temperature compared to those grown at 37ºC (Fig. 7F). These results suggest that enhanced inflammasome formation and caspase-1 activation by NLRC4-H443P mutant at lower temperature is likely to be due to reduced interaction with its negative regulator, HSC70.

**Fig. 7:**
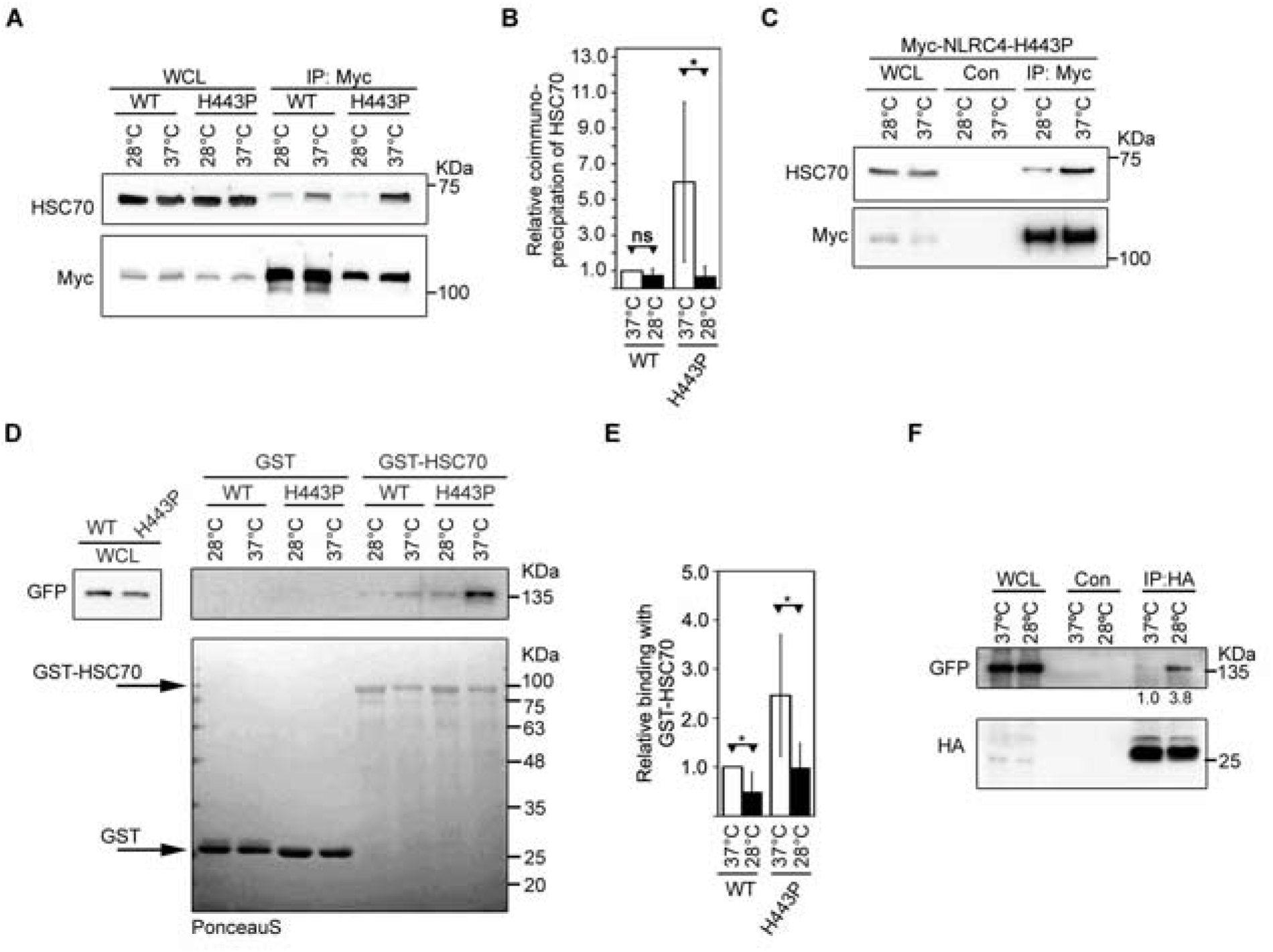
Exposure to subnormal temperature alters interaction of NLRC4-H443P with HSC70 and ASC. **(A)** NLRC4-H443P shows reduced interaction with HSC70upon exposure of cells to subnormal temperature of 28°C for 6h after 18h of transfection. Lysates of HEK293T cells expressing Myc-NLRC4 or Myc-NLRC4-H443P were subjected to immunoprecipitation using agarose conjugated Myc antibody and immunoprecipitates analysed by western blotting. **(B)** Quantitation of relative abundance of endogenous HSC70 in the immunoprecipitates is shown. (n=4); *** p<0.0005. **(C)** Lysates of HEK293T cells expressing Myc-NLRC4-H443P were subjected to immunoprecipitation using agarose conjugated Myc antibody or control antibody, and immunoprecipitates were analysed by western blotting. **(D)** Lysates of HEK293T cells expressing GFP-NLRC4 or GFP-NLRC4-H443P were incubated with GST or GST-HSC70 and bound proteins analysed by western blotting. **(E)** Quantitation of relative abundance of NLRC4 or NLRC4-H443P in pulldown samples of GST-HSC70 is shown. n=3; * p<0.05. **(F)** HEK293T cells were transfected with GFP-NLRC4-H443P along with HA-ASC, and exposed to subnormal temperature of 28°C for 4h after 12h of transfection. Lysates were prepared 16h post-transfection and subjected to immunoprecipitation using agarose conjugated HA-antibody. Western blot analysis shows increase in binding of ASC with NLRC4-H443P upon exposure to subnormal temperature.

## Discussion

NLRC4 is generally activated by pathogen associated molecular patterns (PAMPs)^10^, but certain point mutations result in its constitutive activation, inflammasome formation and caspase-1 activation leading to autoinflammatory diseases^24^. Through this study, we show that HSC70 and HSP70 chaperone proteins form a complex with NLRC4. HSC70 keeps the constitutively active NLRC4-H443P mutant in check to suppress inflammasome formation and caspase-1 activation. Exposure of cells to subnormal temperature results in reduced interaction of HSC70 with H443P mutant, which is accompanied by hyperactivation of caspase-1. Paradoxically, compared to WT-NLRC4, the constitutively active H443P mutant shows enhanced interaction with HSC70. The mutated histidine residue is at a location required for binding with ADP molecule^8^, which is crucial in maintaining inactive configuration of NLRC4. We hypothesize that H443P mutation, in addition to exposing oligomerisation sites by inducing conformational changes in NLRC4, enables stronger interaction of HSC70 and HSP70 with the mutant protein. This mutation possibly induces very specific conformational changes in NLRC4 that are not seen in two other mutants, V341A and T337S, which do not show enhanced interaction with HSC70. Our results also show that regulation of inflammasome activation is dependent on the ATPase activity of HSC70. It is possible that NLRC4-H443P, which shows conformational alterations, is recognized as a misfolded protein by HSC70.

Binding of HSC70 with H443P mutant may be preventing efficient oligomerisation with ASC as shown by increase in speck formation in cells with reduced HSC70 levels. Lower temperature may be inducing a conformational change in HSC70 resulting in reduced interaction with H443P mutant leading to its hyperactivation. This is shown schematically in Fig. 8. This proposed role of HSC70 in mediating low temperature-induced hyperactivation of caspase-1 by H443P mutant is supported by the following observations. Only H443P mutant, which shows enhanced interaction with HSC70, shows enhanced inflammasome formation at lower temperature or upon treatment with HSC70/HSP70 inhibitor apoptozole. The V341A mutant, which does not show enhanced interaction with HSC70, does not respond to lower temperature or apoptozole treatment to form the inflammasome. It may be noted that disease phenotypes of each of the NLRC4 mutants differ and only H443P mutant causes hypersensitivity to low temperature.

**Fig. 8:**
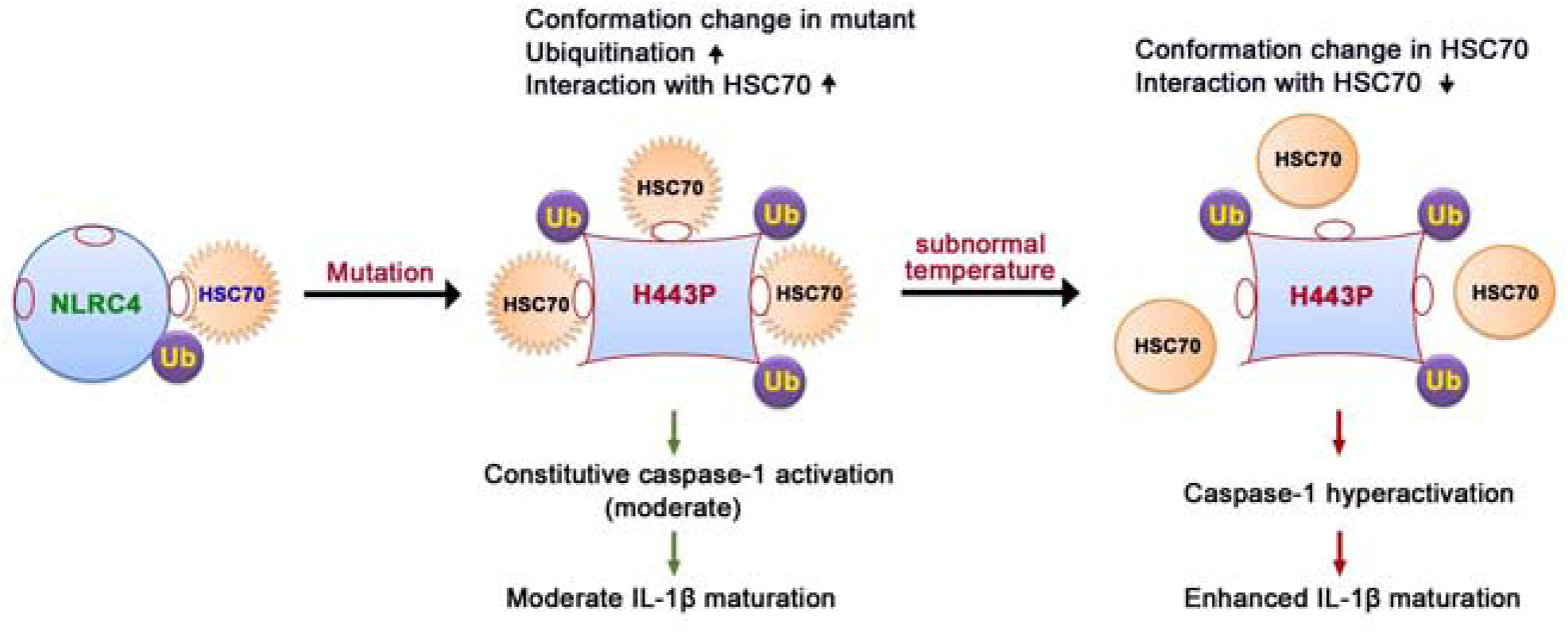
Proposed model for regulation of NLRC4-H443P by HSC70 upon exposure to lower temperature. NLRC4 is present in an inactive closed configuration with low levels of ubiquitination and weak binding to HSC70. Mutation of H443 to proline causes a conformational change in NLRC4, which enables enhanced ubiquitination and more stable interaction with HSC70. H443P mutant shows constitutive caspase-1 activation and moderate IL-1β maturation, causing mild inflammation. Upon exposure of cells to subnormal temperature, HSC70 undergoes a conformational change that lowers its ability to interact with H443P. This allows increased ASC-speck formation by H443P, and caspase-1 hyperactivation leading to enhanced IL-1β maturation and hyper-inflammation. This mechanism of differential interaction of HSC70 with H443P in response to subnormal temperature explains the hyper-inflammation seen in FCAS-patients carrying this mutation.

HSC70 performs several functions in the cell including its role as a chaperone which helps in folding of newly synthesized proteins in the cytosol. The chaperone function involves transient interaction of HSC70 with hydrophobic patches in the native and partially folded/mis-folded proteins^38, 39^. Based on biochemical and biophysical properties it has been suggested that HSC70 may function as a thermal sensor in the physiological temperature range of 30-37°C to adjust the chaperoning activity according to the requirement of the cell^36, 48^. HSC70 undergoes an exothermic reversible transition beginning at about 30°C, and it is more sensitive to protease chymotrypsin at 40°C than at 20°C, indicating a conformational change^48^. These properties of HSC70 support our inference that the reduction of its interaction with H443P mutant seen at lower temperature (28°C) is likely to be due to a conformational change in HSC70.

Previously we have shown that in comparison with WT-NLRC4, H443P mutant undergoes higher level of ubiquitination and interaction with ubiquitinated proteins^35^. In comparison with WT-NLRC4, higher levels of the H443P mutant are seen in cellular complexes formed by HSC70. But, HSC70 did not show enhanced interaction with H443P mutant in GST-pulldown assays unless a deubiquitinase inhibitor was added in the lysis buffer. We suggest that HSC70 interacts with ubiquitinated NLRC4, and higher level of ubiquitination of H443P mutant contributes to enhanced interaction with HSC70. It is likely that ubiquitination induces a conformational change in H443P mutant which results in exposure of HSC70-binding sites. An alternate possibility is that ubiquitination stabilizes an open conformation of H443P mutant with exposed HSC70-binding sites. However, we cannot rule out the possibility of interaction of ubiquitin with HSC70, although the homologous protein HSP70 does not interact with ubiquitin^56^.

H443P mutation in NLRC4 results in enhanced interaction with HSC70 as well as HSP70, and these interactions are drastically reduced at lower temperature (data for HSP70 not shown). Therefore, it is possible that in addition to HSC70, HSP70 may also be involved in mediating low temperature induced increase in inflammasome formation and caspase-1 hyperactivation. However, in comparison with HSC70, the constitutive expression of HSP70 in cells is very low. Furthermore, HSP70 generally mediates its effects upon induction by heat shock. Therefore, the contribution of HSP70 to cold induced hyperactivation of inflammasome formation, if any, is likely to be very small.

We conclude that HSC70 interacts with NLRC4 and H443P mutation alters its conformation to favor a more stable complex with HSC70. Caspase-1 hyperactivation due to H443P mutation is kept in check by HSC70, as its loss or reduction in interaction induced by exposure to cold temperature results in enhanced caspase-1 activation. Our results, therefore provide an understanding of the molecular mechanism for the exacerbation of inflammation induced by cold temperature in individuals carrying the H443P mutation, causative of FCAS.

## Methods

### Cell culture, transfections and treatments

Dulbecco’s modified Eagle Medium (DMEM) containing 10% fetal bovine serum (FBS)was used to maintain HEK293T cells procured from ATCC. RPMI-1640 was supplemented with heat-inactivated 10% FBS and used to maintain human macrophage cell line THP1. Cells were grown at 37ºC in a water-jacketed incubator which maintained 5% CO_2_ and controlled humidity. For transient expression of proteins, plasmids were transfected using Lipofectamine2000 or Lipofectamine3000 (Invitrogen) as per the manufacturer’s protocol. In general, HEK293T cells showed about 80% transfection efficiency.

For siRNA mediated knockdown of HSC70, HEK293T cells were seeded in a 24-well tissue culture plate or on coverslips and transfected with 100 pmols of control siRNA or HSC70-siRNA (sc-29349) for 24h. A second transfection was carried out after 24h, with 100 pmols of control or HSC70-siRNA along with desired plasmids. Lysates were prepared 24h after second transfection for western blot analysis. For immunofluorescence, cells on coverslips were fixed using 4% formaldehyde, 24h after second transfection.

For inhibiting ATPase activity of HSC70, apoptozole was added to the culture medium in desired concentrations, for 12h (unless indicated otherwise), after 12h of transfection with desired plasmids.

For exposure to subnormal temperature, one set of cells expressing desired plasmids were maintained at 37°C, while one set was shifted to 28°C for 6h (unless indicated otherwise) after 18h of transfection.

### Antibodies and Chemicals

Antibodies against GFP (cat no. SC-9996), caspase-1 (SC-56036), myc (SC-40; SC-789), HSC70 (SC-7298), HSP70 (SC-66048), Haemagglutinin (HA; SC-805), IL-1β (SC-7884) and caspase-1 (SC-515) were obtained from Santacruz Biotechnology, USA. Antibodies against cleaved caspase-8 (CST#9496S) and NLRC4 (CST#12421) were from Cell Signaling Technology. Actin (MAB1501) and GAPDH antibodies (MAB-374) were from Millipore.

Agarose conjugated Myc antibody (MycTrap; yta-20), GFP antibody (GFPTrap; gta-20) and control agarose beads (bab-20) used for immunoprecipitation were purchased from ChromoTek, Germany. HRP-conjugated mouse and rabbit secondary antibodies were obtained from GE Healthcare. FemtoLUCENT™ PLUS HRP Kit (786-10) protease inhibitor cocktail was procured from GBiosciences. Cy3 (Indocarbcyanine) conjugated mouse or rabbit secondary antibodies were purchased from EMD Millipore and Alexa633- / Alexa488-conjugated secondary antibodies were procured from Thermo Fisher. NEM (cat no. E3876) was procured from Sigma-Aldrich. Blotto (sc-2324) and apoptozole (sc-221257) were purchased from Santacruz Biotechnology.

### Expression vectors

pEGFP-C1-NLRC4 and its deletion constructs, pCB6-WT-Caspase-1 and pcDNA3-IL-1β have been described previously ^57^.Disease associated mutants GFP tagged NLRC4-H443P, NLRC4-V341A and NLRC4-T337S were generated by CAC-CCC,GTG-GCG and ACC-TCC nucleotide substitutions, respectively. Dr. Barbara Kazmierczak, Yale University kindly provided pCruzMycB-NLRC4 and NLRC4-V341A expression vectors. Myc tagged NLRC4-H443Pand NLRC4-T337S mutants were generated using site directed mutagenesis. cDNAs coding for HSC70 and HSC70-SBD were cloned in BamHI and XhoI sites of pGEX-5X-2 bacterial expression vector (GE Healthcare). Sequences of all the mutants were confirmed by automated DNA sequencing.

### Indirect immunofluorescence and microscopy

Immunostaining of cells fixed with formaldehyde was carried out as described previously^58^. Immunostained cells were observed and images captured with an automated AxioImager Z.2 (Zeiss) fluorescence microscope using Axiovision software under x40/0.75 NA dry objective.

### Quantitation of inflammasome formation by immunofluorescence

HEK293T cells plated on coverslips were transfected with HA-ASC along with Myc tagged NLRC4 and its desired mutants. Exposure to subnormal temperature or treatment with apoptozole was for 6h after 12h of transfection. Cells were fixed 18h post-transfection and immunostained with HA- and Myc-antibodies. In experiments involving siRNA mediated HSC70 knockdown, transfections were carried out as described in the earlier section (*Cell culture, transfections and treatments*) and cells were fixed 48h post-transfection. Immunostained cells were observed under fluorescence microscope and scored for presence of specks in coexpressing cells. Data are presented as mean±S.D. of %cells forming specks from at least 3 independent experiments done in duplicate. At least 300 expressing cells from each coverslip were examined.

### Co-immunoprecipitation and western blotting

For immunoprecipitation of Myc / GFP / HA-tagged proteins expressed in HEK293T cells, agarose conjugated Myc / GFP / HA-antibody was used as described earlier^35^. For immunoprecipitation of endogenous NLRC4, THP1 cells were differentiated into macrophage-like cells by treatment with 10nM PMA for 72h. Differentiated cells were washed with ice-cold PBS followed by lysis in buffer containing 20mM Tris-HCl (pH7.5), 0.5% NP40, 150mM NaCl, 0.5mM EDTA, 1mM PMSF, 0.1% BSA, protease inhibitor cocktail and 10mM N-ethylmaleimide (a deubiquitinase inhibitor). Cells were scraped and collected in a pre-cooled microfuge tube and allowed for lysis at 4°C for 30minutes on a rototorque. Lysate was centrifuged (10000g;10minutes; 4ºC) to remove cellular debris. 2μg of NLRC4 antibody or control IgG was incubated with agarose conjugated Protein A/G for 2h at 4°C before adding to the cell lysates and incubated for 8h at 4°C on a rototorque. The bound proteins were washed thrice with buffer containing 150mM NaCl, 20mM Tris-HCl (pH7.5), 1mM PMSF, 0.5mM EDTA, protease inhibitor cocktail and 10mM NEM. Immunoprecipitates were lysed in SDS containing Laemmli sample buffer. The samples were then subjected to western blot analysis as described earlier^58^. Bands were quantitated using ImageJ software.

### GST-pull down assays

*E. Coli* BL-21 DE-3 cells transformed with plasmid expressing desired GST-tagged protein were induced for protein expression by 1mM isopropyl *β*-D-thiogalactoside (IPTG) for 15h at 18°C. Bacterial cells were lysed by sonication in chilled PBS containing 1mM PMSF and protease inhibitors.1% TritonX-100 was added for solublization and left for 30minutes at 4°C. Lysate was centrifuged for 10minutes at 4°C and 10000*g* to remove insoluble fraction. Glutathione–agarose beads (50%slurry) were added to the supernatant and incubated on a roto-torque at 4°C for 1h to pulldown GST-fusion protein. The beads were pelleted (3000*g*, 4°C, 1minute), washed three times with chilled PBS (containing 1mM PMSF and 0.1% TritonX-100) and incubated with lysates of HEK293T cells (prepared in buffer containing 20mM Tris-HCl (pH7.5), 0.5% NP40, 150mM NaCl, 0.5mM EDTA, 1mM PMSF, 10mM NEM (optional) and protease inhibitor cocktail), transiently expressing desired proteins for 20-30minutesat 37°C, 28°C or 4°Con a roto-torque. Unbound proteins were removed by washing three times at 4°C and bound proteins were boiled in Laemmli sample buffer and analyzed by western blotting.

### Statistical analysis

Quantitative data is represented as mean±S.D. values. Two-tailed Student’s t-Test was used to calculate significance of differences between two means and one-tailed t-Test was used to determine significance of relative difference of a test sample compared to control as 1.

## Acknowledgements

We are grateful to Dr. Barbara Kazmierczak of Yale University, for providing pCruzMycB-NLRC4 and pCruzMycB-NLRC4-V341A expression vectors. This work was carried out with support from Department of Biotechnology, Government of India, awarded to GS and VR (grant no: BT/PR14917/BRB/10/888/2010). GS gratefully acknowledges Department of Science and Technology, Government of India for J.C. Bose National Fellowship (grant no: SR/S2/JCB-41/2010). AKR is grateful to Council for Scientific and Industrial Research, India for fellowship.

## Author contributions

AKR and RR performed the experiments. GS, VR and AKR planned the experiments, analyzed the data and wrote the paper. GS and VR conceived and obtained funding for this study. All authors approved final version of the manuscript.

## Data availability

All data generated or analysed during this study are available from the corresponding author upon reasonable request.

