## SupplementaryFigures for "HSC70 regulates cold-induced caspase-1 hyperactivation by an autoinflammation-causing mutant of cytoplasmic immune receptor NLRC4"

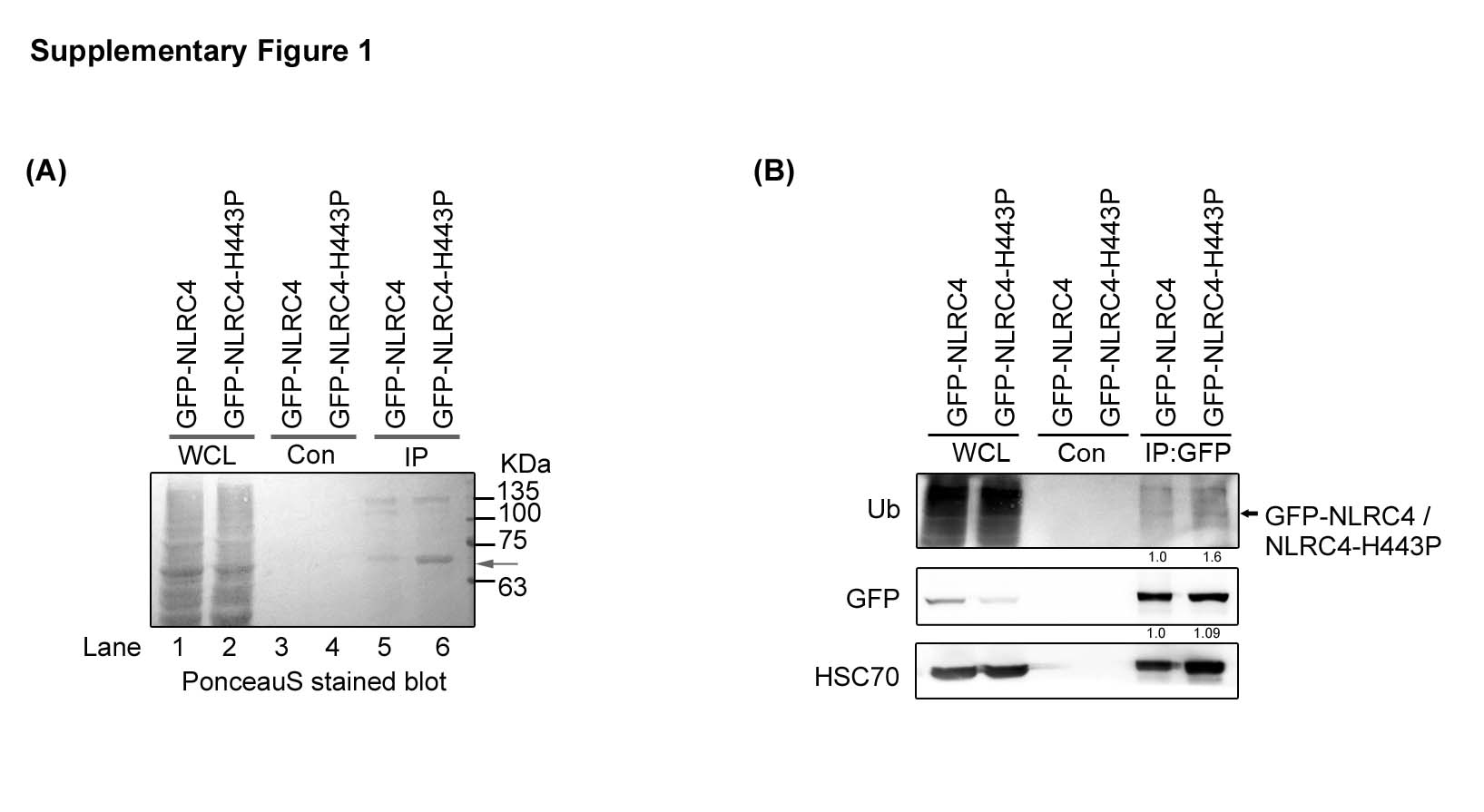

**Supplementary Figure 1**: (A) A ponceauS stained blot of immunoprecipitates of NLRC4 and NLRC4-H443P. Arrow indicates the prominent band of approximately 70KDa. (B) NLRC4-H443P shows enhanced interaction with HSC70 and enhanced ubiquitination. Blot probed with antibodies as indicated. Arrow indicates position of GFP-NLRC4 or GFP-NLRC4-H443P in the blot probed with ubiquitin antibody.

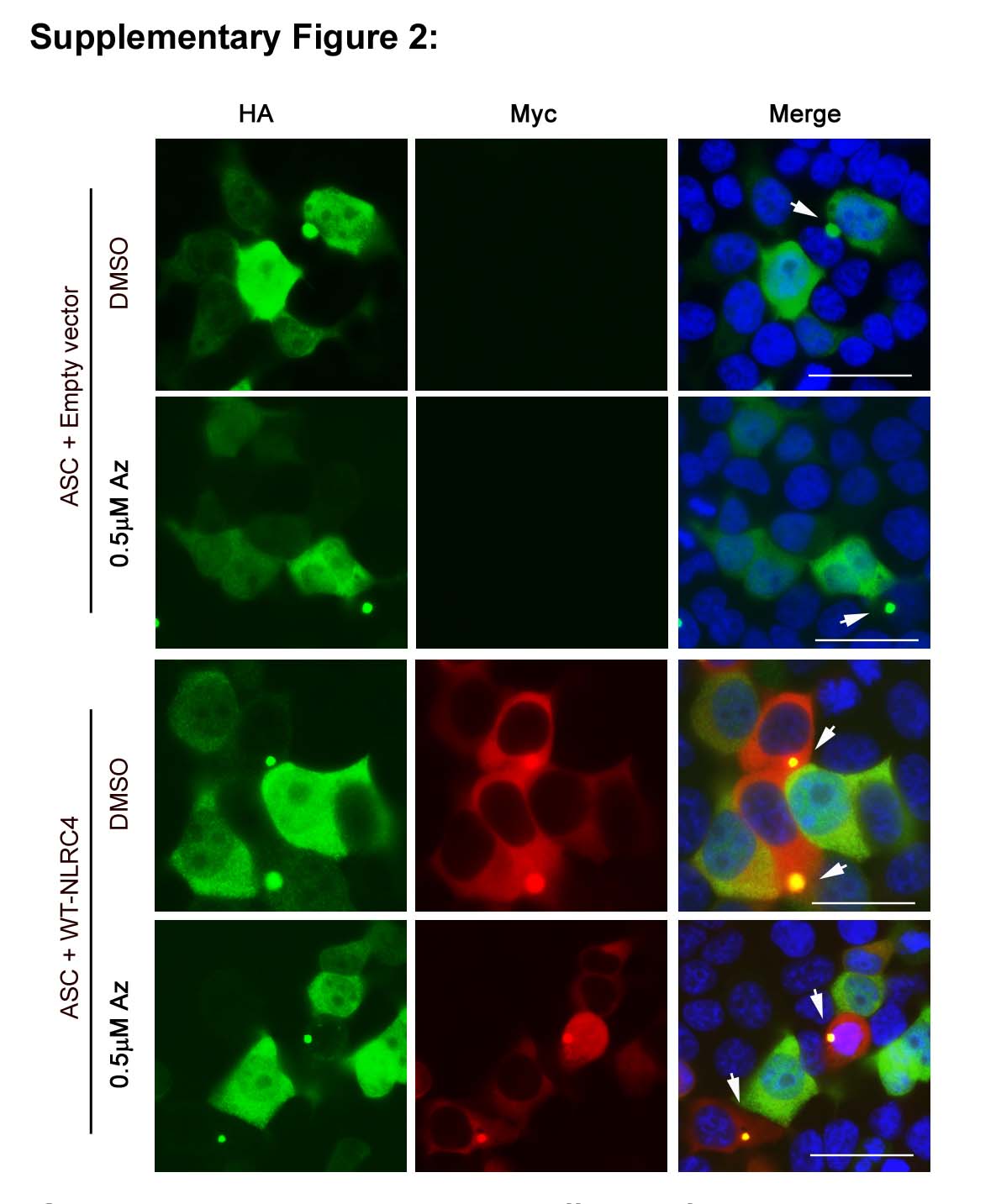

**Supplementary Figure 2**: Effect of apoptozole on ASC-mediated speck formation. Representative immunofluorescence images showing effect of apoptozole treatment on ASC-speck formation by Myc-NLRC4. White arrows indicate specks. DAPI was used to stain nucleus. Scale bars, 20µm.

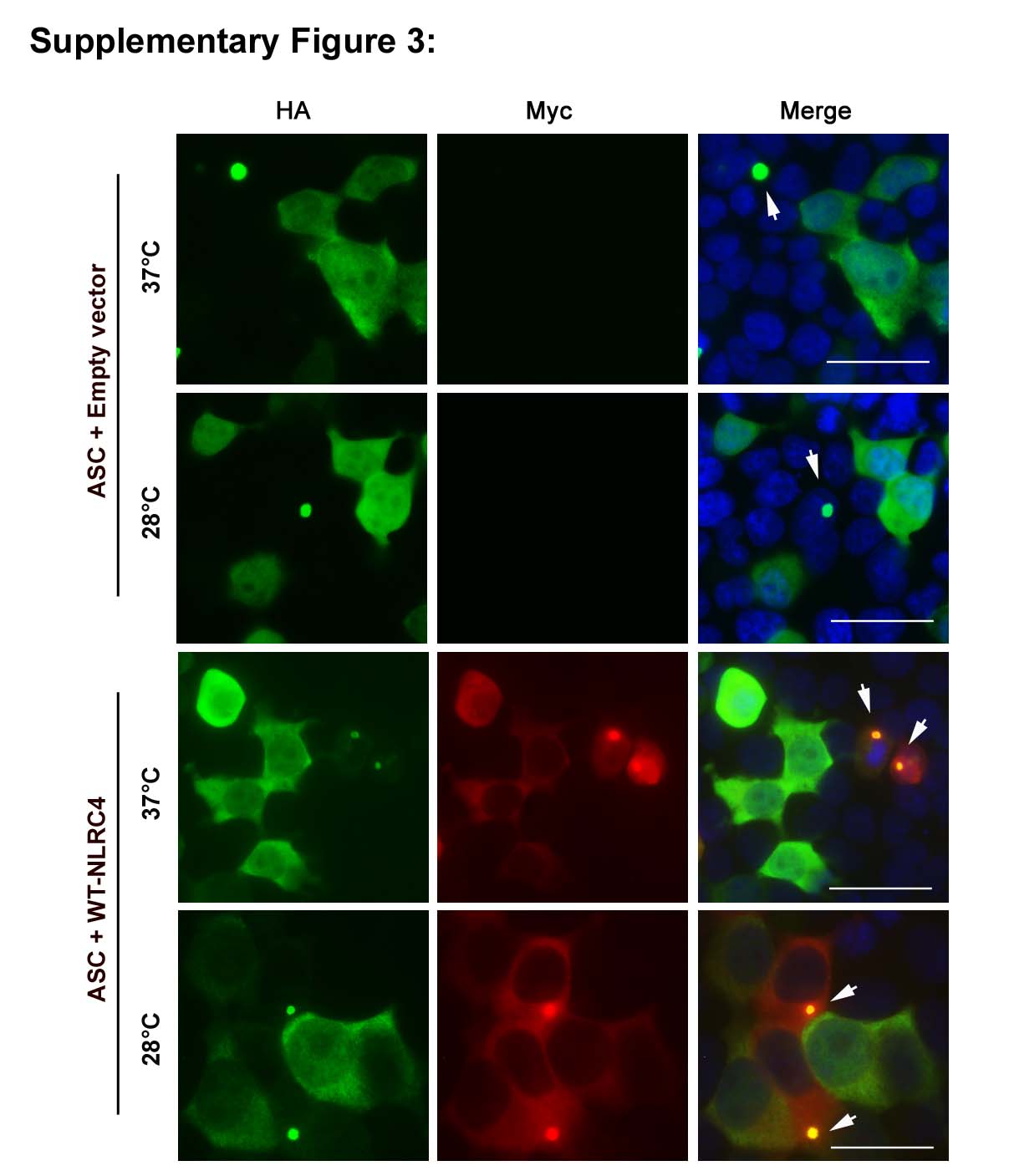

**Supplementary Figure 3:** Effect of exposure to subnormal temperature on ASC-mediated speck formation. Representative immunofluorescence images show effect of exposure to subnormal temperature (28°C for 6h) on ASC-speck formation in cells expressing HA-ASC along with WT-NLRC4. White arrows indicate specks. DAPI was used to stain nucleus. Scale bars, 20µm.
